# Persistent Na^+^ current couples spreading depolarization to seizures in *Scn8a* gain of function mice

**DOI:** 10.1101/2024.10.11.617888

**Authors:** Isamu Aiba, Yao Ning, Jeffrey L. Noebels

## Abstract

Spreading depolarization (SD) is a slowly propagating wave of massive cellular depolarization that transiently impairs the function of affected brain regions. While SD typically arises as an isolated hemispheric event, we previously reported that reducing M-type potassium current (I_KM_) by ablation of *Kcnq2* in forebrain excitatory neurons results in tightly coupled spontaneous bilateral seizure-SD complexes in the awake mouse cortex. Here we find that enhanced persistent Na^+^ current due to gain-of-function (GOF) mutations in *Scn8a* (N1768D/+, hereafter D/+) produces a similar compound cortical excitability phenotype. Chronic DC-band EEG recording detected spontaneous bilateral seizure-SD complexes accompanied by seizures with a profound tonic motor component, which occur predominantly during the light phase and were detected at ages between P33-100. Laser speckle contrast imaging of cerebral blood flow dynamics resolved SD as a bilateral wave of hypoperfusion and subsequent hour-lasting hypoperfusion in *Scn8a*^D/+^ cortex in awake head-restrained mice evoked by a PTZ injection. Subcortical recordings in freely moving mice revealed that approximately half of the spontaneous cortical seizure-SD complexes arose with a concurrent SD-like depolarization in the thalamus and delayed depolarization in the striatum. In contrast, SD-like DC potential shifts were rarely detected in the hippocampus or upper pons. Consistent with the high spontaneous incidence *in vivo*, cortical slices from *Scn8a*^D/+^ mice showed a raised SD susceptibility, and pharmacological inhibition of persistent Na^+^ current (I_NaP_), which is enhanced in *Scn8a*^D/+^ neurons, inhibited SD generation in cortical slices *ex vivo* as well as in head-fixed mice *in vivo*, indicating that I_NaP_ contributes to SD susceptibility. *Ex vivo* Ca^2+^ imaging studies using acute brain slices expressing genetic Ca^2+^ sensor (Thy1-GCAMP6s) demonstrated that pharmacological activation of I_KM_ suppressed Ca^2+^ spikes and SD, whereas an I_KM_ inhibitor strongly increased the frequency of hippocampal Ca^2+^ spikes in *Scn8a*^D/+^, but not WT slices, suggesting that I_KM_ restrains the *Scn8a* GOF hyperexcitability. Together, our study identifies a cortical SD phenotype in *Scn8a* GOF mice shared with the *Kcnq2*-cKO model of developmental epileptic encephalopathy, and reveals that an imbalance of non-inactivating inward and outward tonic membrane currents bidirectionally modulates spatiotemporal SD susceptibility.

## Introduction

Spreading depolarization (SD) is a slowly propagating wave of cellular depolarization of focal origin, involving a profound minute-lasting loss of intra/extracellular ion gradients, massive tissue edema, hypoxia, and transient loss of neuronal activity.^1,2^ Clinical studies reveal frequent SD incidence in traumatic and vascular brain injuries, and SD events can be associated with neurological deficits. In epilepsy cases, peri-ictal and interictal SD may contribute to acute and chronic neurosensory (headache/allodynia, photophobia) and behavioral (depression, anxiety) comorbidities. However, due to the necessity for intracranial recordings for reliable detection, most of the evidence linking SD with these symptoms is available from clinical observation of acute brain injury patients under critical care or experimental models using exogenously provoked SD, often under sedation. The pathological significance of SD in epilepsy patients has yet to be clarified in awake ambulatory settings.

In recent experimental studies using chronic DC-band intracranial EEG recording in a mouse model of *Kcnq2* deficiency linked to developmental epileptic encephalopathy (DEE),^3,4^ we found that conditional deletion in forebrain excitatory neurons (Emx1-Cre: *Kcnq2* flox/flox; hereafter *Kcnq2* cKO) increases tissue SD susceptibility and results in spontaneous bilateral cortical seizure-SD co-generation.^5^ *Kcnq2* encodes the potassium channel pore-forming subunit responsible for the M-type K^+^ current (I_KM_), a non-inactivating slow outward current enriched in the axonal initial segment (AIS).^6,7^ At the AIS, *Kcnq2*/Kv7.2 forms a heterotetramer with *Kcnq3*/*Kv7.3* and regulates local electrogenesis by counteracting depolarizing voltage-gated Na^+^ currents.^8^ This compartmental functional proximity as well as their similar kinetics raises the possibility that increased persistent sodium current (I_NaP_) in excitatory neurons may produce an intrinsic membrane excitability defect equivalent to that seen in *Kcnq2*-cKO mice.

In the mammalian brain, *SCN2A*/Nav1.2 and *SCN8A*/Nav1.6 are the major voltage-gated Na^+^ channel (VGSC) pore-forming subunits expressed in postnatal forebrain excitatory neurons. While both channels are enriched at the AIS,^9^ *SCN8A*/Nav1.6 in particular makes a dominant contribution to I_NaP_.^10–12^ Gain-of-function (GOF) mutations in *SCN8A* associated with enhanced I_NaP_ have been identified in developmental epileptic encephalopathy patients,^13,14^ and a knock-in mouse carrying a heterozygous *Scn8a* N1768D (hereafter *Scn8a*^D/+^)^15^ allele shows enhanced persistent and resurgent Na^+^ current in excitatory neurons^16^, and recapitulates major features of the DEE phenotype including spontaneous seizures, behavioral abnormality, and premature death.

In this study, we characterized SD events in *Scn8a*^D/+^ mice to elucidate the potential role of I_NaP_ in SD regulation. As hypothesized, chronic EEG monitoring of these mice detected spontaneous bilateral seizure-SD complexes, as previously reported in *Kcnq2*-cKO mice.^17^ Furthermore, laser speckle contrast imaging (LSCI) resolved synchronous bilateral propagation of cortical hypoperfusion associated with the SD wave, and subcortical EEG recordings indicated frequent involvement of the thalamus and striatum. Additionally, *ex vivo* assay revealed increased SD susceptibility in acutely prepared *Scn8a*^D/+^ cortical slices which was suppressed by the I_NaP_ inhibitors riluzole and GS967. We also detected possible functional interactions between I_NaP_ and I_KM_ regulating network hyperexcitation. Together, our study reveals an enhanced SD phenotype arising from a clinically identified *Scn8a* GOF mutation and demonstrates that an imbalance of persistent Na^+^ and K^+^ currents produces similar patterns of SD susceptibility in genetic mouse DEE models.

## Material and methods

### Animals

All experiments were conducted under the protocol AN8446 approved by the IACUC of Baylor College of Medicine. *Scn8a*^D/+^ mice were originally generated^15^ and kindly provided by Dr. Miriam Meisler (University of Michigan). These mice were crossed once with the C57BL/6J line at Baylor College of Medicine and the offspring was maintained by inbreeding. In some experiments, *Scn8a*^D/+^ mice were crossed with a Thy1-GCAMP6s GP4.3 mouse (JAX#024275)^18^ for Ca^2+^ imaging studies. All mice were maintained on a 12-hour light-dark cycle with ad libitum access to water and standard chow (5V5M).

### Surgery

Mice received preoperative analgesia (2 mg/kg meloxicam, 1 mg/kg buprenorphine base extended release, s.c.), followed by 2-3% isoflurane anesthesia, and were placed on a stereotaxic frame with a heating pad. The incision site was depilated and cleansed with betadine and 70% ethanol 3 times, and locally injected with a 2% lidocaine/0.5% bupivacaine mixture. The cranial surface was exposed, and 0.5 mm burr holes were made for insertion of a 0.1 mm Teflon insulated silver wire either in the epidural space for cortical surface recording or into subcortical structures, using the following coordinates relative to bregma; cortex (Anterior ±1.0 mm/Lateral: +1.5 mm), hippocampus (Anterior −1.5 mm, Lateral 1.0 mm, depth 2 mm), striatum (Anterior +0.5 mm/Lateral +1.0 mm, depth 2-2.5 mm), thalamus(Anterior −1.5 mm, Lateral 1.0 mm, depth 3.5-4 mm). For implantation in the dorsal pons, bilateral craniotomies (Anterior: −1.0 mm, Lateral: 1.5 mm, relative to lambda) were made to visually identify and avoid blood vessels over the inferior colliculus when inserting the electrode wire. The tips of depth electrodes were marked with DiD fluorescent dye. A ground electrode was placed over the cerebellum through an occipital cranial burr hole. The exact positions of the burr holes were adjusted by ∼1 mm when major blood vessels were present. All wires were connected to an 8-channel pedestal on the skull and cemented with Metabond.

For LSCI studies, the entire dorsal skull was exposed, and epidural electrodes were placed over the somatosensory cortex, olfactory bulb, and cerebellum (ground). A head-bar was placed on the occipital bone. After securing each component using Metabond, the skull surface was coated with cyanoacrylate 3 times.

After the surgery, mice were returned to the vivarium and received postoperative analgesia (2 mg/kg Meloxicam, s.c.) for 3 days, with further recovery for at least 5 days before monitoring. The locations of depth electrodes were verified following recordings by visualizing the fluorescently marked wire tip.

### Chronic EEG monitoring

Mice were transferred to a satellite room where temperature is maintained at 22-24 °C and humidity 40-70% with a 12-hour light-dark cycle. Each mouse was connected to a tethered wire (1 mm diameter) in a recording chamber. DC-EEG signals were amplified with a Bioamp (ADI) along with video using LabChart software (ADI). During monitoring, mice had ad libitum access to water and food, and beddings were replaced weekly.

SDs were first screened based on their voltage and duration thresholds (>5 mV, 30 seconds) and each event was later visually confirmed. A tonic seizure was defined as the chaotic high-frequency EEG activity associated with tonic posturing. In a subset of subcortical recordings, events in one hemisphere were excluded from analysis when depth electrodes were not properly located in the targeted site or the EEG baseline was not stable enough to analyze a DC potential shift.

### Audiogenic seizure

Mice implanted with four cortical EEG leads were transferred to the recording chamber while EEG and motor behavior were recorded. After a ∼20-minute habituation session, a buzzer sound (∼110dB) was delivered for 20 seconds or until a seizure was detected. After monitoring the mice were returned to their homecage.

### Laser speckle contrast imaging

Mice were maintained on a treadwheel using a metal bar head restraint. The dorsal surface of the skull was illuminated with an 800 nm LED (1-2 mW) through a series of lenses to enhance the speckle pattern, and images acquired with a CMOS camera (DMK 33UX273) fitted with an IR-pass filter and polarizer. A temporal contrast image was generated from fifteen serially captured images acquired at 50 Hz using MATLAB at 1 Hz. The contrast values were used as a measurement of cerebral blood flow (CBF). EEG and EMG signals were obtained (Bioamp). A thermistor (36AWG, 5TC-TT-K-36-36-ROHS, Omega) was used to capture nasal airflow. Overall movement was monitored using a standard CMOS camera through an IR cut filter.

SD was provoked by an intraperitoneal injection of pentylenetetrazol (PTZ) at a subconvulsive dose determined in pilot studies using separate cohorts (titrated from 20 mg/kg in *Scn8a*^D/+^ and 40 mg/kg in WT). SD responses in some mice were examined repetitively, in which case each injection was separated by 7-10 days to minimize any kindling effect.

### *Ex vivo* SD studies using acute brain slices

Acute slices were prepared using methods described previously^17^ and details are provided in the supplementary document. During recording, slices were transferred to a submerged chamber (RC-27, Warner Instruments) continuously perfused with ACSF at 2.5 mm/min and maintained at 33-34°C. Both intrinsic optical signal (IOS) and fluorescence were acquired with a CMOS camera (Hamamatsu) controlled with µManager software (https://micro-manager.org/). Field potentials were acquired with glass micropipettes filled with ACSF, amplified by a MultiClamp 700B, and digitized (Digidata 1550B, Molecular devices).

In the KCl SD threshold test, the IOS was acquired at 0.5 Hz while coronal cortical slices were exposed to ACSF containing incrementally elevated KCl concentrations, starting with 6 mM, and incremented by 1 mM every 5 minutes until an SD was detected, as in our previous study.^17^

In the Ca^2+^ imaging study, GCAMP6s fluorescence from horizontal slices was excited using a fluorescence light source (Lambda-DG4, Sutter) and acquired for 30 ms at 1 or 2 Hz. Spontaneous seizure-like activity and SD were induced by exposure to nominally Mg^2+^ free ACSF (0 mM MgSO_4_) described in our previous study^19^. The raw fluorescence images were converted to ΔF/F_0_ and analyzed using ImageJ software. Ca^2+^ activities corresponding to seizure-like activities and SD were initially defined based on their correlation with the field potential and were later determined based on the intensity, duration, and migration speed of the Ca^2+^ activity. The Ca^2+^ spike frequency was determined based on the number of Ca^2+^ spikes above the threshold (3 times the standard deviation) during exposure to Mg^2+^-free solution excluding the duration of SD.

### Drugs

ICA 110381, Riluzole, and XE991 were purchased from Tocris, and GS967 from AOBIOUS. In *ex vivo* studies, drugs were dissolved in DMSO at 100 mM, added to ACSF, and dissolved by sonication. All other chemicals were purchased from Sigma. In the *in vivo* study, 100 mM GS967 solution was suspended in 1% methylcellulose and intraperitoneally injected.

### Statistics

All data analyses are performed using Labchart, MATLAB, pClamp10, R, and prism software. Two group comparisons were performed by a *t*-test, while multigroup comparisons were conducted by ANOVA and post-hoc Sidak test. The results of ANOVA are described in figure legends, and the result of post-hoc tests are in the figure panels when p-values <0.05. In bar graphs, data are presented as mean ± standard deviation with individual data points. Experiments were blindly performed where it is possible.

### Data availability

Data supporting the findings of this study are available within the article and its Supplementary material. All supporting data in this study are available from the corresponding author on request.

## Results

### A bilateral seizure-SD complex is the predominant EEG abnormality in *Scn8a*^D/+^ mice

We characterized the cortical SD phenotype in a total eleven *Scn8a*^D/+^ mice (4 males and 4 females *Scn8a*^D/+^ mice and three male *Scn8a*^D/+^:Thy1GCAMP6s) with chronic DC-band EEG recordings starting from the age of P33-80. Each mouse was implanted with four cortical surface electrodes (**Figure 1A**). During the initial week of recording, almost all generalized seizures were followed by a large negative DC potential shift nearly simultaneously detected at all four electrodes (**Figure 1B**), similar to an EEG event frequently seen in Emx1-*Kcnq2* cKO mice.^17^ The DC potential shifts at the anterior and posterior electrodes were similar in amplitude (14.9 ± 7.5 mV vs 16.0 ± 8.6 mV, p=0.4, n=71) and duration (58.3 ± 28.9 s vs 63.4 ± 24.3 s, p=0.26, n=71).

**Figure 1.**
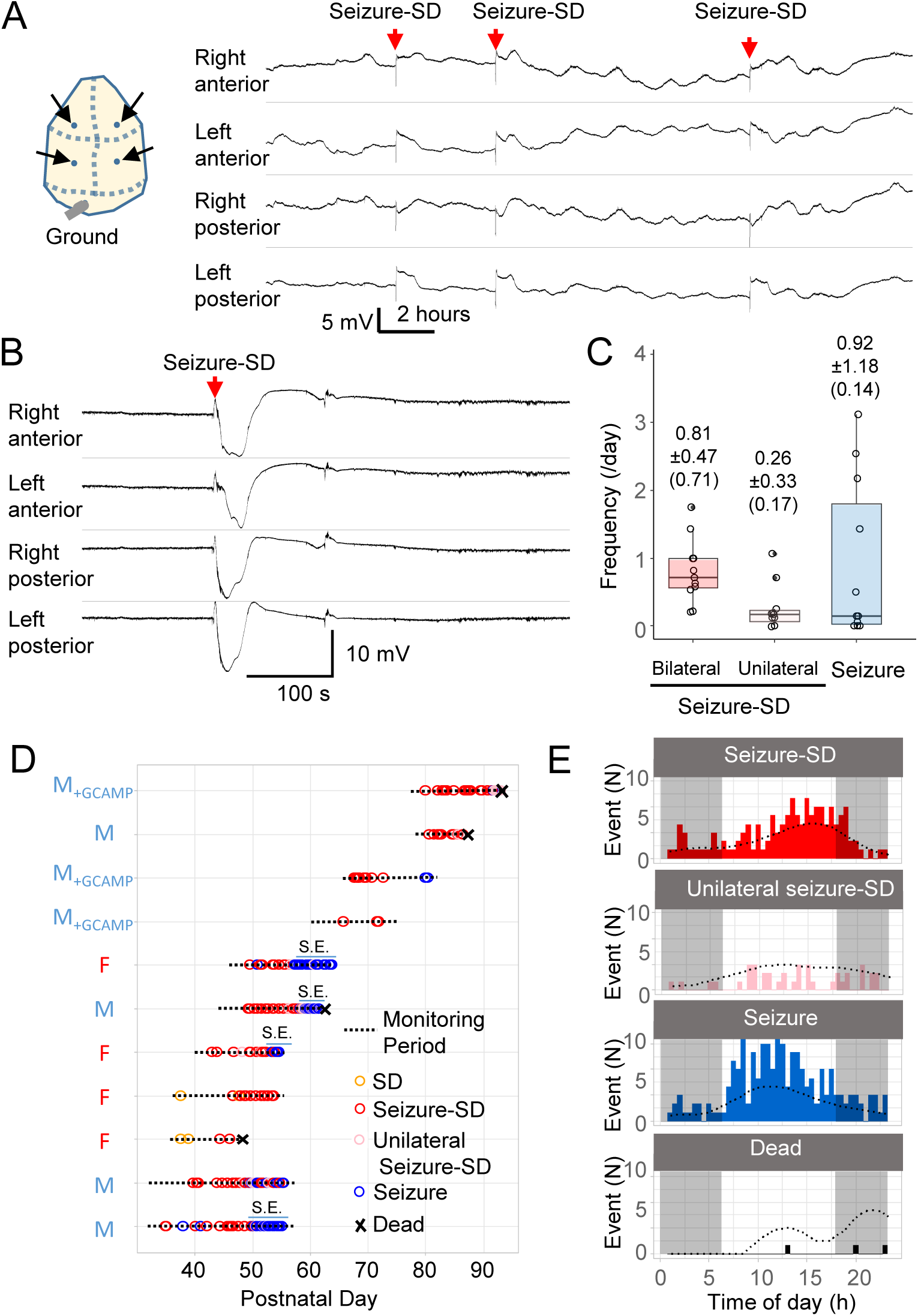
Chronic DC-band EEG recording in *Scn8a*^D/+^ mice. **A.** Scheme showing cortical EEG electrode implant positions and an example of a ∼12-hour EEG trace. **B.** Expanded EEG trace of bilateral seizure-SD complex. **C.** Box-whisker plots showing the frequencies of bilateral seizure-SD complexes (left), unilateral seizure-SD complexes (middle), and seizures (right). The values over the plots indicate mean standard ± deviation and median values in parentheses. **D.** Raster plot of cortical EEG events across ages in the *Scn8a*^D/+^ cohort. M: male, F: female **F.** Cumulative histograms of detected cortical EEG events across time of day. The gray shade indicates dark phases, and the dashed lines show a density plot.

Bilateral seizure-SD complexes and solitary seizures were the major EEG events whose mean frequencies were less than once per day (**Figure 1C**), however, these events tended to appear as a cluster and multiple events could be detected in a single day (see **Figure 1A**). During the ensuing weeks of serial recording, five mice started to show irregular DC potential shifts and/or SDs detected only unilaterally (**Figure 1D**, Unilateral Seizure-SD). In these mice, the DC potential shift eventually became undetectable, leaving only a mostly electrographic seizure signature (**Figure 1D**, Seizure) which tended to involve less robust motor manifestations. In four of these mice, seizure frequency increased and eventually evolved into status epilepticus (by definition recurrent seizures with less than an hour interictal interval, **Figure 1D**). One of the mice did not recover and died (**Supplementary Figure 1**), while another required euthanasia.

Both seizure-SD complexes and seizures showed a diurnal pattern; these events were more frequent during the light phase than in the dark phase with slightly distinct peaks (seizure-SD peak at 15.9, seizure peak at 11.4, **Figure 1E**).

### Seizure-SD complexes were accompanied by tonic seizures

Each seizure-SD complex was always associated with a generalized motor seizure featuring a robust tonic component characterized by a hunched back with spastic extension of both fore- and hind limbs as reported in the original characterization of this transgenic mouse.^15^ An example of the seizure-SD complex is presented in **Figure 2**. In EEG recordings, the ictal seizure phase was detected as chaotic high-frequency activity, often contaminated by muscular fibrillation artifact, and preceded or followed by slower sharp wave seizure activity associated with clonus (**Figure 2B**). These pre-SD tonic seizures were always present during the onset of the DC-potential shift. The slow baseline drift and the subsequent sharp spikes seen in the high-pass filtered trace (**Figure 2B**) are nearly identical to the electrographic seizure characterized using an AC-coupled amplifier,^15^ reinforcing the necessity of unfiltered DC amplification for reliable detection of SD. The peak of the DC potential shift was followed by a minutes-lasting depression of EEG amplitude (**Figure 2A**), while it could be interrupted by one or two additional post-SD tonic seizures detected in 66.6% of events (**Figure 2D**).

**Figure 2.**
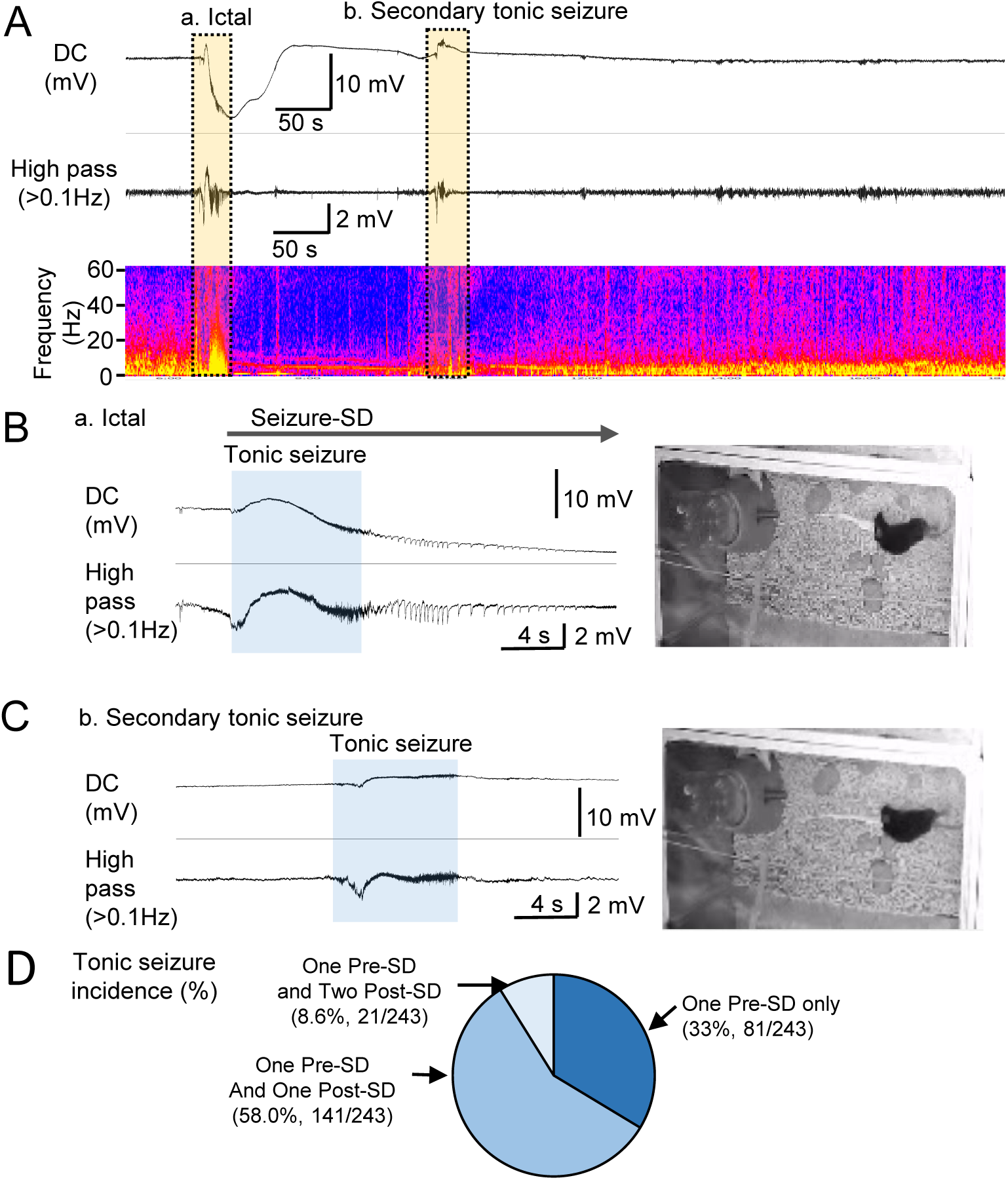
Electrographic characterization of cortical seizure-SD complex in *Scn8a*^D/+^ mice. **A.** Representative DC-band and high-pass filtered (>1Hz) EEG traces and power spectrum of a seizure-SD complex. The expanded EEG traces in orange boxes corresponding to a. ictal and b. tonic seizure are presented in **B&C**. SD onset was preceded by a seizure characterized by a robust tonic seizure detected as high-frequency noise (blue shade), followed by slow ictal spikes (**B**). After an SD, another tonic seizure was detected as high-frequency activity (blue shade) (**C**). Mouse posture associated with tonic seizures is typically characterized by a hunched back and hind limb extension as shown in the images at right. **D.** The number of tonic seizures during a seizure-SD episode. About one-third of events were associated with only one tonic seizure at the SD onset, while the rest of the seizure-SD episodes were associated with one or two additional post-SD tonic seizures.

Tonic seizures were rarely detected in the absence of electrographic seizure or seizure-SD complex during chronic monitoring. In a few instances, we detected a DC shift that preceded the tonic seizure, suggesting that seizure and SD may not always propagate as a complex, or that an EEG seizure is not a necessary trigger of SD in this model.

### Tonic spasm during audiogenic seizure does not involve cortical SD

Next, we examined whether cortical SD is present during the tonic spasm during an audiogenic seizure in *Scn8a*^D/+^ mice. In awake young *Scn8a^D/+^* mice (P30-45), a buzzer sound provoked audiogenic seizures, characterized by wild running followed by sudden motor arrest in an extended tonic spastic posture (**Figure 3**, upper). All recorded mice spontaneously recovered from the audiogenic seizure and none died. During the tonic phase, we detected a chaotic high-frequency EEG activity similar to that seen during a spontaneous tonic seizure associated with a seizure-SD complex. In all six recordings, audiogenic seizures appeared without the robust cortical DC potential shifts suggestive of SD (**Figure 3**, lower). Thus, while chronic monitoring revealed the frequent appearance of tonic seizures with cortical SD, tonic seizures can be triggered as an isolated paroxysmal event without a cortical SD.

**Figure 3.**
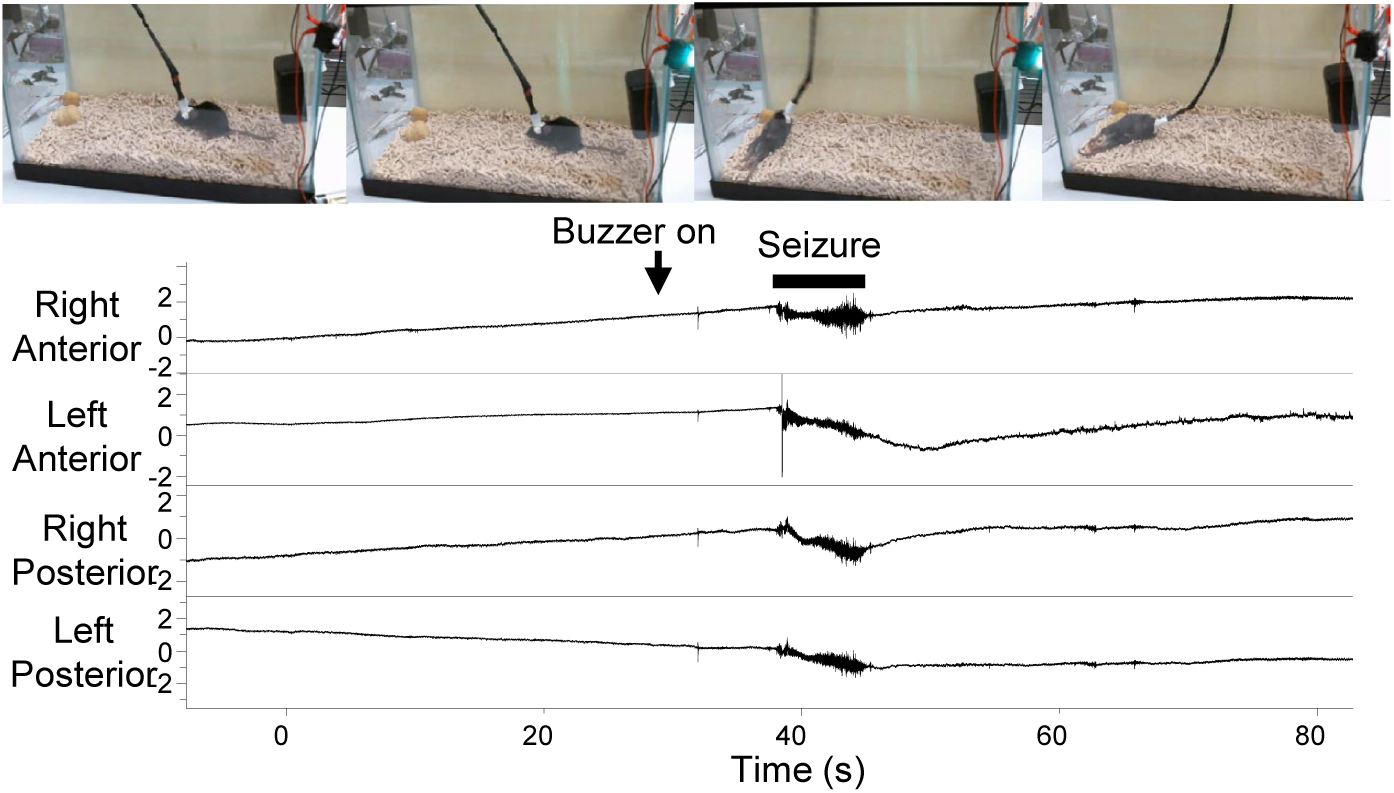
Audiogenic tonic motor seizure did not involve cortical SD. An audiogenic seizure was provoked by a loud buzzer in *Scn8a*^D/+^ mice implanted with cortical EEG electrodes as shown in **Figure 1**. A robust tonic spasm was detected during audiogenic seizure, however, no SD-like DC potential shift was detected in all six recordings.

### Bilateral SD wave detection by laser speckle contrast imaging

In order to gain further insight into the pathophysiology of the bilateral SD event, we conducted a laser speckle contrast imaging (LSCI) in head-restrained awake mice to characterize the spatiotemporal profile of SD based on the CBF dynamics.^20^ No spontaneous seizure-SD complexes were detected in a total of >30 hours of monitoring from eight *Scn8a*^D/+^ mice. Therefore, we utilized a PTZ injection reported to trigger bilateral SD in non-epileptic awake rats.^21^ In *Scn8a*^D/+^ mice, a PTZ injection (20-30 mg/kg, i.p.) triggered a bilateral slow DC potential shift (16.1 ± 3.0 mV, 79.8 ± 29.4, s, n=8) in 62% (8/13) of trials in five *Scn8a*^D/+^ mice (**Figure 4A&B**). Similar to spontaneous seizure-SD complexes, these SDs were preceded by a brief seizure and visualized as a bilaterally-synchronous wave of cerebral hypoperfusion which always appeared at the lateral edge of the dorsal cortical surface and spread toward the midline (**Figure 4A, Video2**). The wave of hypoperfusion reaches the midline within 15-20 seconds after its appearance, suggesting that SD spread through the cortex at a rate >15 mm/minute. This exceptionally fast speed may be due to sustained elevation of K^+^ during the prior seizure, and/or partial inhibition of GABA_A_Rs by PTZ. The initial wave of hypoperfusion was followed by a transient partial recovery and subsequently by an hour-lasting hypoperfusion (**Figure 4B**). Tonic seizures associated with SD were temporally associated with enhanced EMG signal, respiratory arrest (apnea), and abrupt decrease in cerebral blood flow (CBF) (**Figure 4C**). These neurological effects were often followed by massive salivation (**Figure 4C&D**), which often attenuated the nasal airflow signals. After recovery from seizure-SD, the mouse typically loses the grip on the wheel along with prolonged immobility (**Figure 4D**, postictal).

**Figure 4.**
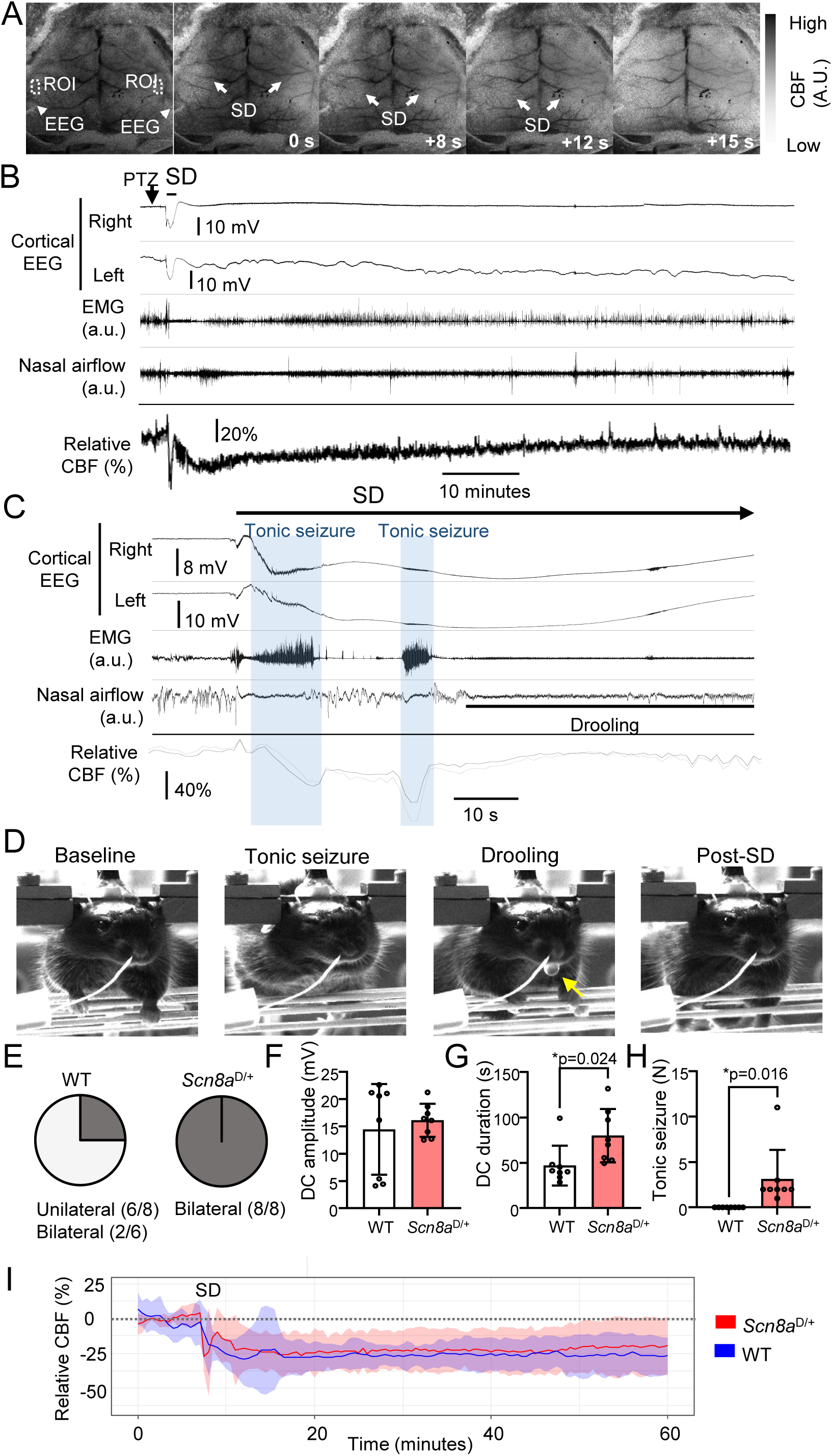
Combined electrophysiological and laser speckle contrast imaging of head-restrained awake mice injected with PTZ. **A.** Sequence LSCI images of bilateral SD detected as waves of hypoperfusion (wavefront shown with white arrow) advancing symmetrically from the lateral to the midline. The raw contrast value (standard deviation/mean) from the region of interest (ROI) was used as the CBF readout. **B.** Cortical EEG, EMG, nasal airflow, and relative CBF changes during an episode of seizure-SD complex after a single PTZ injection. The onset of cortical SD with tonic seizures is expanded in **C**. Tonic seizures (blue shade) were detected as enhanced EMG signal at the onset and during cortical SD. During tonic seizures, nasal airflow was absent and CBF was slightly reduced. **D.** Images of mouse behavior. During baseline, the mouse is firmly grabbing the running wheel to maintain posture. During tonic seizure, forelimbs are retracted and eyes are partially closed. After the second tonic seizure, massive salivation was detected. After SD, the mouse became immobile and often did not grab the running wheel. **E.** SD waves in *Scn8a*^D/+^ mice were always bilateral, while only 25% of them were bilateral in WT. The amplitudes of the DC potential shift were similar (**F**), while the duration was prolonged in *Scn8a*^D/+^ mice (**G**). **H.** Tonic seizures were only seen in *Scn8a*^D/+^ mice. **I.** The mean trace of normalized CBF changes in WT and *Scn8a*^D/+^ mice from 5 recordings. Shading indicates 95% confidential interval.

In the control experiments using WT mice, a higher dose of PTZ (40-60 mg/kg) was necessary to trigger an SD event. Epileptic activities were also variable in WT mice, preceded by a brief generalized convulsive and/or recurrent myoclonic spikes which often outlasted SD. SDs in the majority of WT mice were unilateral (75%, 6/8, **Figure 4E**, Fisher’s exact test, p=0.007 vs. *Scn8a*^D/+^). SD detected in WT mice had a similar amplitude (**Figure 4F**, 14.4 ± 8.3 mV, n=8, p=0.46), while the duration of DC potential shift was ∼40% shorter than in *Scn8a*^D/+^ (**Figure 4G**, 46.9 ± 22s, n=8, p=0.024). None of the SDs in WT mice were associated with tonic seizure (**Figure 4H**), indicating that the paroxysmal tonic spasm associated with SD was a specific feature of *Scn8a*^D/+^ mice. Similar to *Scn8a*^D/+^ mice, SDs in WT were detected as a wave of hypoperfusion originating at the lateral cortical convexity and spreading toward the midline. CBF responses in the minutes after SD passage were variable in WT mice, likely reflecting variable post-SD epileptic activities, whereas SD affected cortex invariably underwent chronic hypoperfusion similarly to *Scn8a*^D/+^ (**Figure 4I**).

Together, these results indicate that the *Scn8a* ^D/+^ mouse cortex is intrinsically susceptible to bilateral SD *in vivo*. To the extent that the PTZ-evoked SD reproduced spontaneous events, the bilateral spreading pattern and a fast propagation rate could explain the near-simultaneous appearance of DC potential shift in all four electrodes during chronic monitoring (**Figure 1B**). The propagating pattern also suggests that the bilateral SD wave likely originates from either somatosensory barrel cortex, or even from more distant regions, such as the piriform cortex and amygdala^22^ (see Discussion).

### Subcortical involvement of seizure-SD complex

Elevated SD susceptibility is associated with SD invasion into subcortical regions.^23–25^ Since the tonic phase of an audiogenic motor seizure is likely mediated by subcortical and brainstem motor pathways, we conducted additional chronic monitoring to examine the subcortical involvement of seizure-SD complex.

#### 1) Hippocampus

SD in the hippocampus was not common in *Scn8a*^D/+^ mice. In hippocampal recordings, depth electrodes were positioned within the CA1 region or in the dentate gyrus (DG). Slow DC shifts in CA1 stratum radiatum (Amplitudes 20.7 ± 3.0 mV, Duration: 49.9 ± 13.0 s) were detected only in a mouse during cortical seizure-SD complexes, and no robust DC potential shifts were detected in the remaining four mice (**Figure 5A**). Overall, intrahippocampal DC shifts temporally associated with cortical seizure-SD complex were detected in 17.5% (11/63, N=5 mice) and were relatively rare (but see Discussion). Tonic motor seizures were evident in the hippocampal recordings as a high-frequency activity similar to those detected in the neocortex.

**Figure 5.**
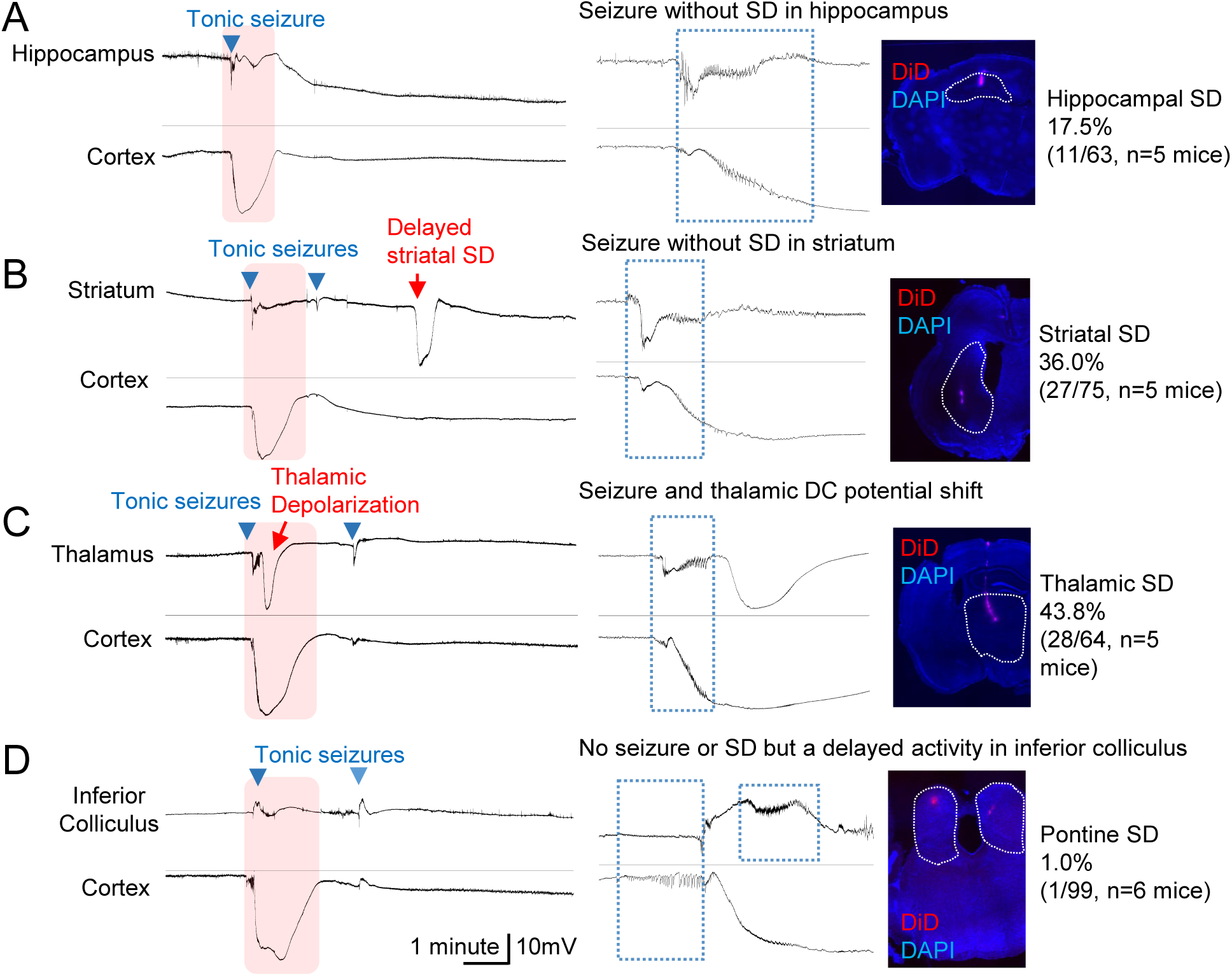
Combined cortical surface and subcortical EEG recordings in *Scn8a*^D/+^ mice. **A.** Cortical seizure-SD complex simultaneously recorded in cortical and hippocampal electrodes. Electrographic **s**eizures were simultaneously detected both in the cortex and hippocampus, however, SD was often absent in the hippocampus. The histology image on the right shows the electrode recording site determined by DiD fluorescence marking the electrode tip. **B.** In striatum recordings, cortical seizure-SD complex was detected as seizure only, however, a delayed DC potential shift was often detected. **C.** In the thalamus, a sharp DC potential shift was often detected in association with a coincident DC potential shift in the cortex. Tonic seizures were detected as relatively large DC potential shifts in this region. **D.** Recordings from inferior colliculus and the rostral pontine nucleus. In these structures, cortical seizure-like discharge was absent, while delayed activity was detected during cortical SD. SD was rarely detected. In this structure, tonic seizures were detected with variable patterns, either with or without DC potential shift. Blue arrowheads indicate tonic seizure, red shade indicates the duration of cortical SD, and red arrows indicate DC potentials suggestive of SD in the subcortical traces. In the middle column, the dashed blue boxes show seizures or abnormal fast activities.

#### 2) Striatum

Delayed SD-like DC potentials were detected in the striatum in association with 36.0% (27/75, N= 5 mice) of cortical seizure-SD complexes. These SD-like DC potential shifts (amplitude: 11.3 ± 6.6 mV, duration: 30.2 ± 13.2 s) appeared after a 144.8 ± 97 s delay (**Figure 5B**). Similar delayed DC potential shifts have been reported, and are considered to reflect a cortical SD wave invading the striatum.^26^ Tonic seizures were also detected in the striatum as high-frequency activity.

#### 3) Thalamus

Recordings from ventrobasal thalamic nuclei detected large DC potential shifts coincident with 47.8% (28/64, N=5 mice) of cortical seizure-SD complexes (**Figure 5C**). These thalamic depolarizations were large amplitude but shorter-lasting (amplitude: 9.6 ± 4.6 mV, duration: 36.0 ± 20.0 s) than the concurrent cortical DC shift. Tonic motor seizures were detected in the thalamus as a negative DC potential shift with overriding high-frequency activity.

#### 4) Inferior colliculus and rostral pontine nucleus

SD-like DC potentials were rarely detected in the inferior colliculus (n=4 mice) and pontine reticular nucleus (n=2 mice). Robust synchronized ictal discharges were absent in both structures, while unique 2-3 Hz activities were detected in both regions during cortical SD (delayed activity in **Figure 5D**). A DC potential shift in the inferior colliculus was detected only once in a mouse undergoing a moribund decline prior to death (**Supplementary Figure 2**). At these sites, tonic seizures were variably detected either as a high-frequency activity with or without DC potential shift either in a positive or negative direction.

Together these subcortical recordings demonstrate DC potential shifts in the SD susceptible striatum and thalamus, while they are rare in the hippocampus and dorsal pons. DC potential shifts in these subcortical regions never preceded a cortical event, and therefore represent either secondary invasion of a cortical SD or independent locally generated depolarization.

### I_NaP_ contributes to SD threshold *ex vivo and in vivo*

We next examined the effect of *Scn8a* GOF mutation on intrinsic tissue SD susceptibility using acute cortical slices prepared from adult *Scn8a*^D/+^ and littermate WT mice. SD susceptibility was measured based on the step-wise incremental elevation of bath KCl, and in this method, the KCl concentration that triggers an SD is used as a threshold (**Figure 6A**, see Methods). This assay detected a small decrease in the KCl threshold in *Scn8a*^D/+^ cortical tissue (**Figure 6B**).

**Figure 6.**
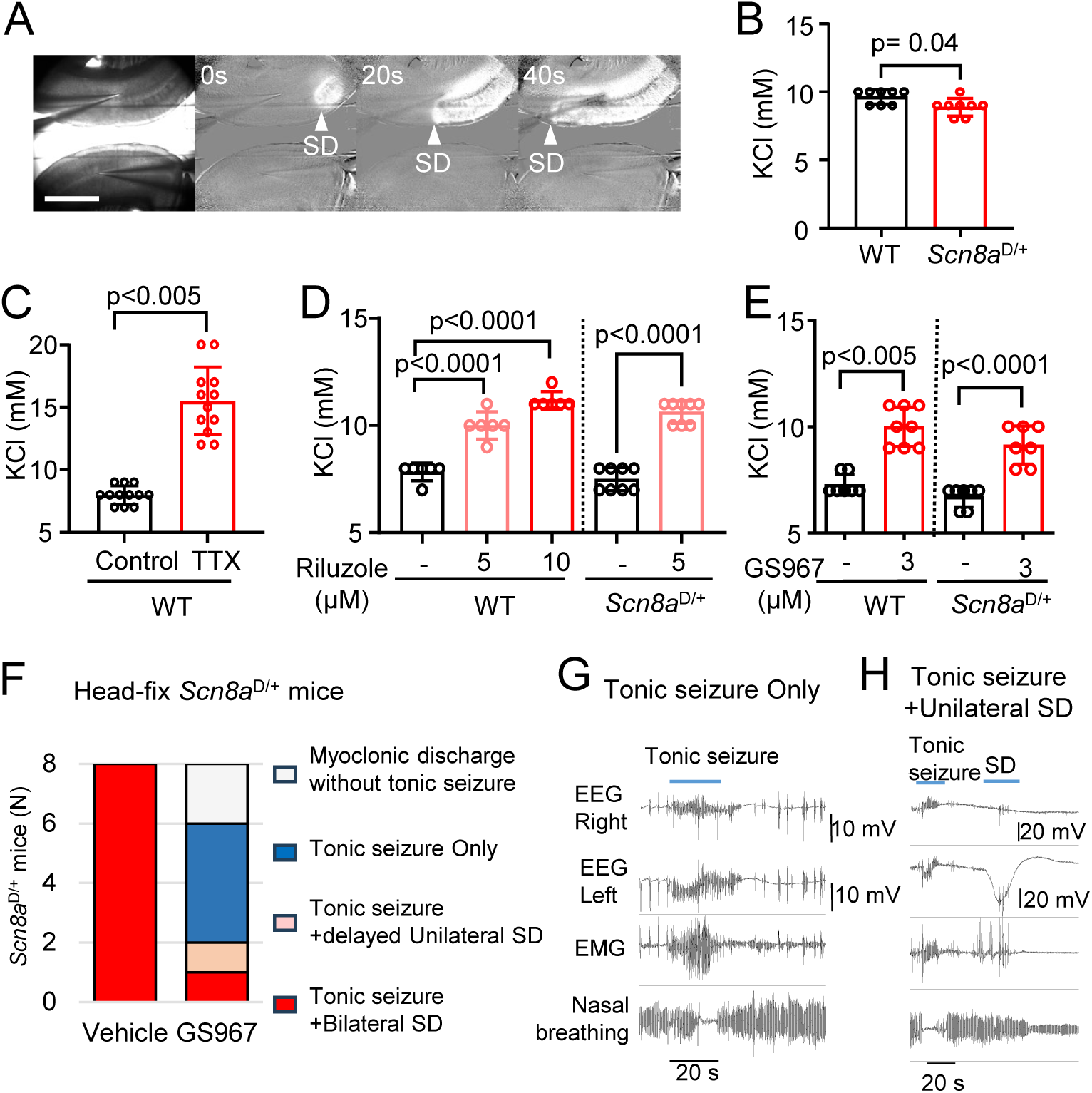
*Ex vivo* SD threshold test revealed a contribution of I_NaP_ to the SD threshold. Slices were incubated in ACSF while KCl concentration was incrementally elevated by 1 mM until SD was generated. **A.** Raw image (left) and ratio images (2^nd^-4^th^ images). An SD wave can be readily detected as an enhanced IOS signal. Arrows mark the leading edge of SD. Scale bar 500 µm. **B.** The bath KCl concentration sufficient to trigger an SD was slightly lower in *Scn8a*^D/+^ slices. **C.** TTX increased the KCl concentration required to trigger SD. N=12 slices. **D.** I_NaP_ inhibitor riluzole increased the KCl concentration to trigger SD in WT and *Scn8a*^D/+^ slices. In WT, One-Way ANOVA, p= <0.0001. N=6 slices each. *Scn8a*^D/+^, N=8 slices each. **E.** A similar inhibitory effect was seen with the I_NaP_ inhibitor GS967. From left, N= 7, 8, 7, 7 slices. **F-H** GS967 inhibits SD in head-fixed *Scn8a*^D/+^ mice *in vivo*. Head-fixed *Scn8a*^D/+^ mice were pretreated with vehicle or GS967 (1.5mg/kg, i.p.) 15 minutes before PTZ injection (20 mg/kg, i.p.). The fraction of responses to PTZ is presented in the bar graphs (**F**), and representative EEG, EMG, and nasal breathing traces of GS967 treated mice with tonic seizure only (**G**) and tonic seizure with unilateral SD (**H**) are presented. N=8 mice each.

The similarity of the cortical bilateral EEG phenotype seen in *Scn8a*^D/+^ and I_KM_ deficient *Kcnq2*-cKO mice raises the possibility that non-inactivating I_NaP_ could directly contribute to the enhanced SD susceptibility. I_NaP_ inhibition decreases SD susceptibility in anesthetized mouse cortex,^27^ however, the role of I_NaP_ in the intrinsic SD threshold *ex vivo* has not been well established. Thus, we analyzed the effect of the I_NaP_ inhibitors Riluzole and GS967^28^ on the SD threshold using the same *ex vivo* SD model.

We first examined the effect of TTX, which non-selectively blocks voltage-gated Na^+^ currents (VGSC). The addition of 1 µM TTX to the bath solution robustly increased the KCl threshold for triggering SD (**Figure 6C**), validating the previously reported^29–32^ contribution of VGSCs in SD generation in our assay system.

We then tested the effect of I_NaP_ inhibitors. Riluzole inhibits I_NaP_ with an IC_50_ in the 2-4 µM range.^33,34^ We used Riluzole at 5 and 10 µM, which could inhibit 75-95% of I_NaP_^33^, 10-32% of I_NaT_^35,36^, and 10-20% of glutamate release evoked by high extracellular K^+^.^37^ Riluzole at both concentrations increased the KCl threshold for SD in both WT and *Scn8a*^D/+^ cortical slices (**Figure 6D**). A second potent I_NaP_ inhibitor, GS967 at 3 µM, a dosage that preferentially inhibits I_NaP_ over transient Na^+^ current,^38,39^ also increased KCl threshold for SD in both WT and *Scn8a*^D/+^ cortical tissue (**Figure 6E**). In this comparison, a *Scn8a* genotype difference was not detected because of insufficient statistical power to detect the small mutation effect.

We next examined the effect of I_NaP_ inhibition by GS967 (1.5mg/kg), previously shown to reduce spontaneous seizures in *Scn8a*^D/+^ mice,^40^ on the susceptibility to seizure-SD complexes in head-fixed *Scn8a*^D/+^ mice. All vehicle-treated mice (1% methylcellulose) developed bilateral seizure-SD complex after PTZ injection (100%, 8/8 mice, **Figure 6F**). In contrast, SD was detected in 25% (2/8) of the GS967 pretreated mice (p=0.007 vs. vehicle, Fisher’s exact test), and the rest of the mice showed myoclonic spikes and/or tonic seizures only (**Figure 6G**). In a GS967 treated mice SD was unilateral and appeared 33 seconds after a tonic seizure (**Figure 6H**). GS967 did not change the incidence of tonic seizure (vehicle 100% (8/8), GS967 75% (6/8), p=0.2, Fisher’s exact test). Overall, GS967 showed a potent inhibitory effect on SD and uncoupled it from seizure *in vivo*.

These results demonstrate that I_NaP_ can influence SD susceptibility *ex vivo* and *in vivo*, while neuronal discharges mediated by transient Na^+^ currents likely have an additional downstream effect of I_NaP_ during the initiation of SD.

### I_KM_ regulates *Scn8a*^D/+^ GOF mediated hyperexcitability

The appearance of SD closely associated with seizure activity in ambulatory mice suggests that SD is generated as a result of hyperexcitable cortical activity. To assess this relationship, we performed *ex vivo* Ca^2+^ imaging using acute horizontal brain slices obtained from *Scn8a*^D/+^ and WT mice carrying a Thy1-GCAMP6s transgene in which a Ca^2+^ sensor GCAMP6s is widely expressed in various brain regions.^18^ Our low-magnification fluorescence imaging detected basal Ca^2+^ signals in somatosensory and entorhinal cortices as well as in the hippocampus, and exposure to magnesium ion-free bath solution resulted in the generation of large spontaneous fast Ca^2+^ spikes during seizure-like field potentials as well as SD detected as a slowly migrating Ca^2+^ wave (**Figure 7A&B**).^19,41^ While such Ca^2+^ signals were occasionally detected in the striatum and thalamus, they were not reproducible and not analyzed here.

**Figure 7.**
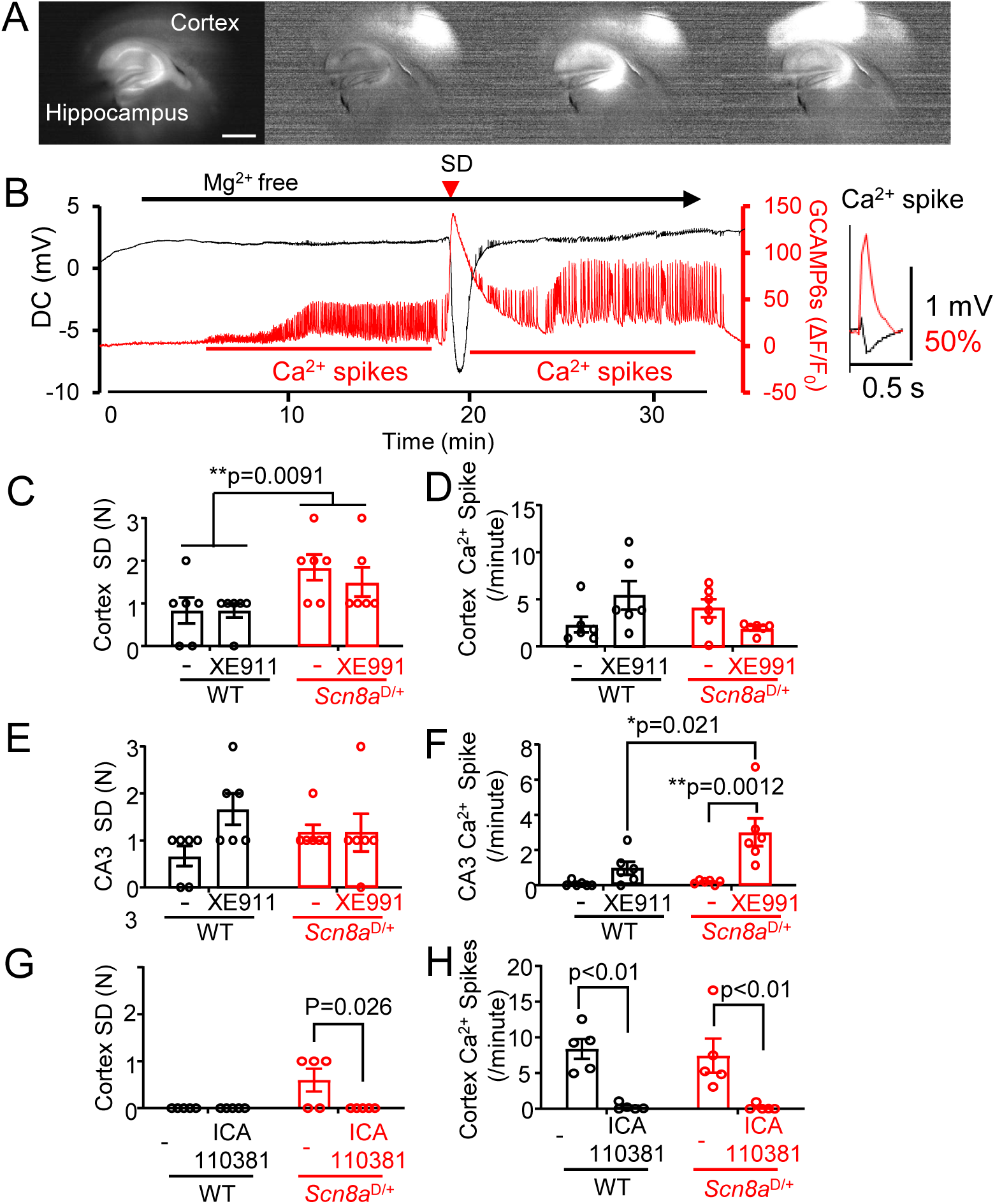
*Ex vivo* Ca^2+^ imaging analysis of the effect of I_KM_ modulators on cortical and hippocampal SD and epileptic activities. **A&B.** Representative images and traces of tissue excitability test by *ex vivo* Ca^2+^ imaging. Horizontal slices obtained from WT-or *Scn8a*^D/+^-GCAMP6s slices were incubated in Mg^2+^ free ACSF. **A.** Raw and ratio fluorescence images showing Ca^2+^ activities in the cortex and hippocampus CA3. SD was detected as a prolonged Ca^2+^ elevation whereas seizure-like activities were detected as fast Ca^2+^ spike frequencies (**B** inset). Scale bar 500 µm. **C-F** Effect of I_KM_ inhibitor XE991. **C.** Cortical SD were more frequently generated in the *Scn8a*^D/+^ than WT and XE991 was without effect (genotype: p=0.009, XE991: p>0.57, interaction: p>0.57) **D.** There was no genotype or XE991 effect on cortical Ca^2+^ spike frequencies (genotype: p>0.41, XE991: p>0.60, interaction: p=0.017). **E.** The SD frequency in the hippocampus CA3 was not modified by genotype or XE991 treatment. (genotype: p>0.10, XE991: p>0.99, interaction: p>0.10). **F.** Spike frequency in the hippocampal CA3 region was low in the control condition, but XE991 greatly enhanced it in *Scn8a*^D/+^ slices (genotype: p=0.024, XE991: p=0.0004, interaction: p=0.039). Statistics were calculated by two-way ANOVA with post hoc Sidak test and the *p*-values of post hoc tests are shown in the corresponding figure panel when they reach the significance level. **G&H** The effect of I_KM_ activator ICA-110381 (3 µM) was tested in a separate cohort. **G.** In these experiments, SD was detected only in the *Scn8a*^D/+^ cortex and was eliminated by ICA-110381. (genotype: p=0.026, ICA-110381: p=0.026, interaction: p=0.026) **J**. Ca^2+^ spikes in WT and *Scn8a*^D/+^ cortex were also eliminated by ICA-110381. (genotype: p=0.76, ICA-110381: p<0.0001, interaction: p=0.72). Statistics were calculated by two-way ANOVA with post hoc Sidak test and the p-values are shown in the corresponding figure panel when they reach the significance level.

In the somatosensory cortex, the number of SD events generated was higher in *Scn8a*^D/+^ than in the WT cortical slices (**Figure 7C**), demonstrating the higher tissue SD susceptibility in *Scn8a*^D/+^ cortical tissue. In contrast, the number of Ca^2+^ spikes was not different between genotypes (**Figure 7D**). Bath application of the M-current inhibitor XE991 (10 µM) did not influence the SD or spike frequency in the cortex (**Figure 7C&D)**. In the hippocampus, SD was most consistently detected in the CA3 region and often remained confined within this subfield in both *Scn8a*^D/+^ and WT slices, therefore analysis focused on the CA3 subfield. Neither the *Scn8a* genotype nor XE911 affected the frequency of hippocampal SDs (**Figure 7E**). Unlike the cortex, Ca^2+^ spikes in the CA3 were less frequent in the control condition, however, I_KM_ inhibition by XE991 greatly enhanced them in *Scn8a*^D/+^, but not in the WT slices (**Figure 7F**), suggesting that basal I_KM_ activity restrains *Scn8a* GOF hyperexcitability in this circuit.

In a separate cohort, we next examined the effects of the I_KM_ activator ICA-110381, which preferentially potentiates *Kcnq2/3*-mediated current with an EC_50_ = 0.38 µM.^42^ Because of the uncertainty in pharmacokinetics, I_KM_ pharmacology study was limited to *ex vivo*. These experiments detected fewer Ca^2+^ activities than the experiment above and since SD and spikes in the CA3 were absent in most slices, analysis was focused on cortical events. ICA-110381 at 3 µM eliminated cortical SD (**Figure 7G**) and greatly reduced the number of Ca^2+^ spikes in both WT and *Scn8a*^D/+^ tissue (**Figure 7H**). These results demonstrate that I_KM_ enhancement effectively suppresses cortical hyperexcitability and suggest that the absence of I_KM_ inhibitor XE991 effects seen in the cortical tissue (**Figure 7C&D**) likely reflects low basal I_KM_.

Together, these results demonstrate a higher SD susceptibility in the *Scn8a*^D/+^ cortex under hyperexcitable conditions. In addition, the robust increase in hippocampal spiking in the *Scn8a*^D/+^ hippocampal tissue indicates that I_KM_ activity counteracts the *Scn8a*^D/+^ GOF mutation effect.

## Discussion

The present study characterized the spatiotemporal seizure-SD phenotype of *Scn8a*^D/+^ mice, revealing a bihemispheric seizure-SD complex as a signature cortical EEG abnormality in this sodium channel GOF mouse model. LSCI imaging and subcortical EEG recordings studies suggest that the seizure-SD complexes are spontaneously generated in the somatosensory or ventral cortex, and about half of the depolarizing waves spread into the striatum and thalamus, and rarely into hippocampus and brainstem. The similar bilateral seizure-SD phenotype seen in *Kcnq2*-cKO mouse cortex suggests that these shared phenotypes can be attributed to an imbalance in the persistent cation currents (I_NaP_, I_KM_). Our results collectively indicate that these specific excitability defects can significantly modulate initiation of the self-propagating wave of slow depolarization and suggest that pharmacologically targeting these cation currents might be useful for controlling the SD threshold in epileptic brain.

### Regulation of SD threshold by persistent Na^+^ and M-type K^+^ currents

The present study identifies *Scn8a* as an SD modifier gene. Given the effect of the N1768D GOF mutation on late sodium channel inactivation, as well as the inhibition of SD by I_NaP_ blockers, our study suggests that I_NaP_ is a significant component contributing to SD susceptibility. SD inhibition by the I_NaP_ inhibitors riluzole and GS967 is consistent with their effectiveness at normalizing the *Scn8a* GOF effect on neuronal defect in early afterdepolarization,^43^ seizures, and premature death in *Scn8a*^D/+^ mice,^40^ as well as reducing seizures in *SCN8A* GOF patients.^44^ Our findings are also in line with a previous study reporting that veratridine, a VGSC opener that functionally mimics the *Scn8a*^D/+^ mutation effect,^45^ induces seizure-SD complexes in acute hippocampal slices, and I_NaP_ inhibitors reduce them.^46^

The shared bilateral cortical seizure-SD phenotype seen in *Scn8a*-GOF and Emx-*Kcnq2* cKO mouse models likely reflects a mutually antagonizing effect of I_NaP_ and I_KM_. Both I_NaP_ and I_KM_ have slow inactivation kinetics and are activated near the resting membrane potential (∼-65 mV).^47,48^ Previous studies demonstrated their functional interaction as inhibition of I_KM_ results in the I_NaP_-mediated burst discharges^49–52^ or a plateau depolarization when extracellular Ca^2+^ is reduced.^53,54^ The latter might be particularly relevant to the slow ramp membrane depolarization during SD which involves a rapid drop of extracellular Ca^2+^.^1^

On the other hand, since *Scn8a* and *Kcnq2* are expressed differentially across cell types and brain regions, mutations in these genes will variably impair the I_NaP_ and I_KM_ balance. For example, *Scn8a*^D/+^ mutation increases the excitability of hippocampal pyramidal neurons in both CA1 and CA3 regions, whereas it selectively enhances interneuron excitability in CA3, but not in CA1.^43^ *Kcnq2* GOF mutations increase the excitability of cortical pyramidal neurons but dampen the excitability of hippocampal pyramidal neurons.^55^ Similarly, I_KM_ mediated by *Kcnq* channels is critical for the medium afterhyperpolarization in CA1 pyramidal neurons,^49,56^ but SK-type channels may play a larger role in cortical pyramidal and other cell types.^57–59^ Such cell type-specific effects may underlie the complex sensitivity to the I_KM_ inhibitor XE991 seen in our *ex vivo* studies (**Figure 7**).

In addition to excitatory neurons, a recent study suggests that increased I_NaP_ in inhibitory neurons due to a *SCN1A* gene mutation identified in familial hemiplegic migraine type 3 (FHM3)^60^ can contribute to SD susceptibility.^61^ This interneuronal mechanism might also contribute to SD susceptibility in the *Scn8a*^D/+^ mice, especially in the CA3 circuit where the GOF mutation enhances I_NaP_ in interneurons.^43^

Both Nav1.6/*Scn8a* and Kv7.2/*Kcnq2* channels are highly enriched at the AIS. Correlatively, this compartment has a lower activation threshold for I_NaP_ ^62^ compared to the soma^51,63^ and dendrites,^64,65^ and I_KM_ density is higher at the AIS than in the soma.^66,67^ Thus, the AIS is likely a vulnerable region to I_NaP_/I_KM_ imbalance due to genetic mutation, and a hyperexcitable AIS may contribute to SD susceptibility by increasing action potential frequency.

In addition to the AIS, I_NaP_ and I_KM_ coexist in other subcellular compartments relevant to SD. One is the axonal terminal boutons where Nav1.6/*Scn8a*^68^ and *Kcnq* channels are detected in some synapses,^69–72^ and regulate transmitter release. In fact, inhibition of Na^+^ current at the presynaptic release site was considered as a mechanism by which the I_NaP_ inhibitor riluzole inhibits glutamate release.^37^

Studies also suggest I_NaP_ and I_KM_ could modify postsynaptic currents. Immuno-electron microscopy studies have detected Nav1.6/*Scn8a*^68^ and Kv7.2/*Kcnq2*^73^ in dendritic spines. In support of a postsynaptic role of I_KM_. several I_KM_ activators act as functional NMDAR antagonists, likely by stabilizing the Mg^2+^ channel block,^74,75^ which would contribute to their SD inhibitory effect.^17,76^ While the physiological significance, molecular composition, and cell/species-specificity are not fully understood, postsynaptic I_NaP_/I_KM_ dysfunction may influence SD susceptibility.

### SD and tonic seizures

Profound tonic spastic seizure without or minimal clonus is a robust clinical phenotype in *Scn8a*^D/+^ mice. *Scn8a* is expressed in murine spinal motoneurons,^77^ and the GOF mutation may directly increase susceptibility to neurogenic spasms.^78^ The frequent appearance of tonic spasms during and after seizure-SD complex suggests that SD might create an excitatory brain state favoring this paroxysmal motor event. On the other hand, unlike other models,^79^ SD was absent when tonic spasm was provoked by an audiogenic seizure, indicating that cortical SD is not required to evoke this response.

Recurrent tonic seizures accompanied by bilateral SD could be reproduced in *Scn8a*^D/+^ mice injected with PTZ (**Figure 4**). These tonic seizures were associated with abrupt apnea, previously associated with diaphragm spasm^80^, as well as a rapid global CBF decrease and subsequent massive drooling (**Figure 4**). We speculate the latter autonomic responses reflect parasympathetic vasovagal pathway activation which could reduce cerebral perfusion pressure and enhance salivation. SD was always followed by an hour-lasting period of hypoperfusion, which was previously detected with a laser-doppler system during KCl-induced SD in properly perfused brain,^81^ and a similar prolonged hypoperfusion occurs after a hippocampal seizure.^82^ These hour-lasting hypoperfusion are mediated by COX1/2 metabolites,^82–84^ and the pattern seen in our model likely shares the underlying molecular mechanism.

Tonic seizures are also a prominent clinical feature of *KCNQ2*-associated developmental encephalopathy patients,^85,86^ suggesting a pathophysiological similarity to *SCN8A* encephalopathy. However, our previous studies on mice with forebrain-specific *Kcnq2* deletion (Emx1-*Kcnq2* cKO) did not detect robust tonic seizures as seen in *Scn8a*^D/+^ mice. This might reflect localization of the *Kcnq2*-sensitive spastic mechanism outside the forebrain such as in the extrapyramidal pathways^87^ and/or spinal motoneurons,^88,89^ or the *KCNQ2*-related clinical phenotype is not fully recapitulated in the mouse.

### SD and seizures in subcortical structures

Our subcortical recordings revealed remote depolarization of subcortical regions associated with cortical seizure-SD complexes. In the striatum, cortical SD was followed by delayed depolarization. A previous study suggested this is the invasion of a cortical wave spreading through the piriform/amygdala complex.^90^ If the propagation pattern is shared, SD invasion into these ventral cortical areas may contribute to autonomic dysfunction in *Scn8a*^D/^+ mice.

We detected SD-like DC potential shifts in the thalamus. Thalamic SD has been reported in a *Canca1a* GOF model and was accompanied by transient hypertension^23^. The thalamic DC potential shift detected in the *Scn8a*^D/+^ mice closely coincided with cortical depolarization and was therefore unlikely to represent an invasion of the cortical wave. Rather, the near-simultaneous cortico-thalamic depolarization suggests it is related to the putative synaptic activation of thalamic neurons during cortical SD wave as detected in awake mice.^91^

SD incidence was rare in the hippocampus. This result was somewhat unexpected because hippocampus, especially CA1, is relatively SD-susceptible,^1^ and the excitability of pyramidal neurons in the hippocampus and adjacent entorhinal cortex is increased in *Scn8a*^D/+^ mice.^43,92^ SD threshold in the hippocampus might be less sensitive to I_NaP_/I_KM_ imbalance or, alternatively, this could be related to SD resistance reported in highly epileptic tissue.^93^ On the other hand, we cannot exclude the possibility that our recording with a wire electrode could have missed a localized SD event as was seen in our *ex vivo* Ca^2+^ imaging study.

The inferior colliculus and rostral pontine nucleus rarely show SD events, which is consistent with their low SD susceptibility, and a heterozygous *Scn8a*^D/+^ mutation did not overcome the high SD threshold. However, SD susceptibility in these brainstem structures might be increased in younger animals with less myelination^94–96^ and a higher susceptibility to abnormal excitation as suggested by their higher susceptibility to audiogenic seizures. In some monogenic epilepsy models, SD in these hindbrain structures are related to sudden death,^96,97^ but such a study was not possible in *Scn8a*^D/+^ model due to their low spontaneous mortality. These brainstem structures lacked ictal discharges detected in the cortex, yet displayed clear activity associated with tonic motor seizures, suggesting that brainstem circuits at this level are involved in the paroxysmal spastic pathology. Fine mapping of the brainstem circuitry involved in tonic seizures may facilitate the development of therapeutics relevant to brainstem hyperexcitability associated with *SCN8A* and *KCNQ2* mutations.

In summary, the present study identified a shared susceptibility to bilateral SD wave generation in *Scn8a* GOF and *Kcnq2*-cKO mice, which suggests a putative role of I_NaP_ and I_KM_ balance in the regulation of SD generation and propagation. The frequent interaction between seizures and SD seen in these DEE models suggests that SD may contribute to the spectrum of comorbidities in patients associated with these ion channelopathies.

## Competing interests

None

### Supplementary materials

Supplementary material is available at *Brain* online

Video 1

Mouse behaviors associated with a seizure-SD complex described in Figure 2.

Video 2

CBF response during seizure-SD complex analyzed in LSCI.

---

**Supplementary Figure 1.**
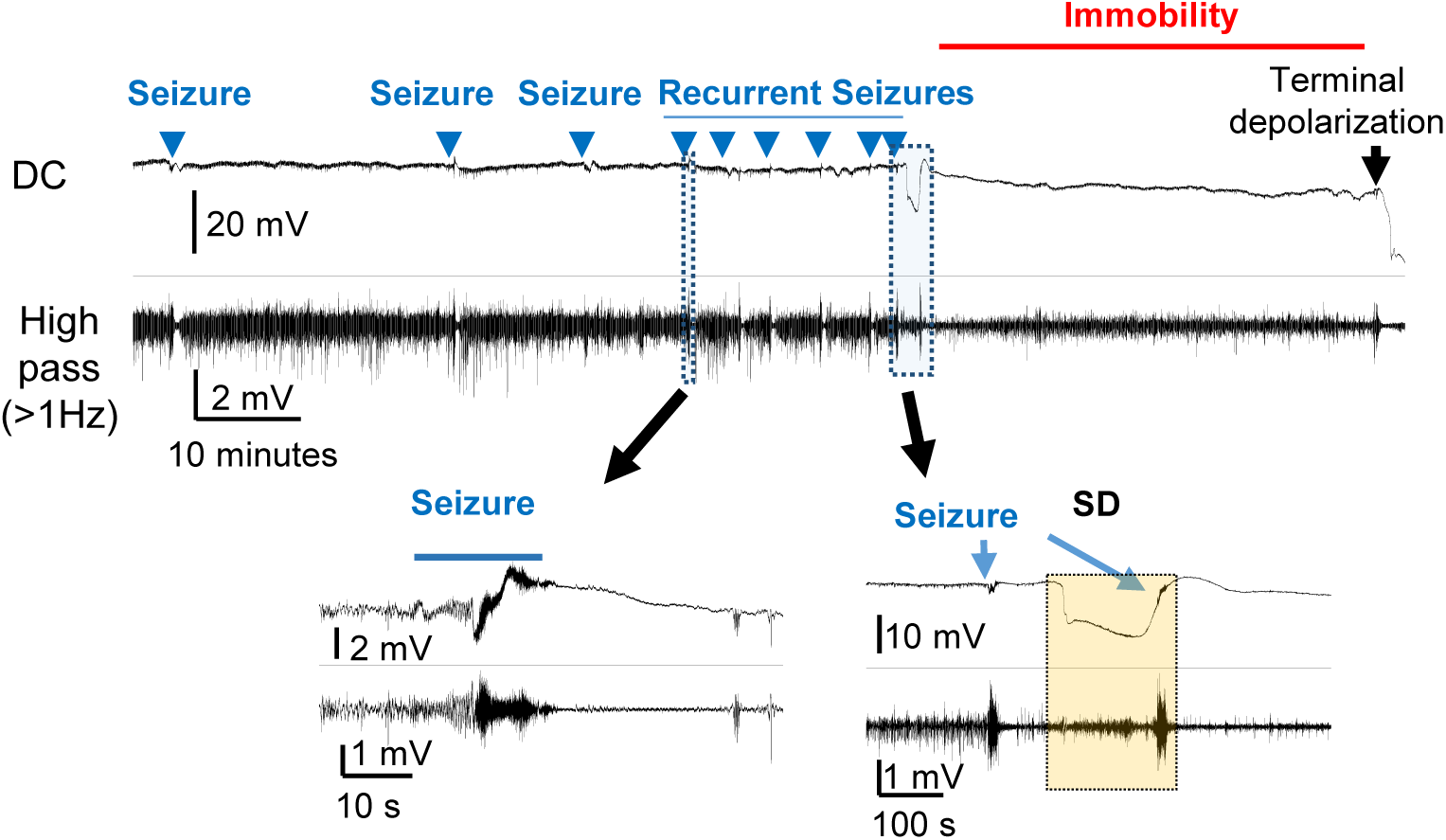
A *Scn8a*^D/+^ mouse that underwent recurrent seizures or status epilepticus. Cortical EEG trace of *^Sc^n8a*^D/+^ mouse which underwent status epilepticus. During status epilepticus, each seizure did not involve overt behavioral convulsions such as wild-running, fall, or spasm. The last seizure was detected during the recovery phase of SD and the mouse underwent prolonged immobility and eventually died as indicated by terminal depolarization. Top: raw DC-band, bottom: 1Hz high-pass filtered trace. Expanded traces of a single seizure and a seizure-SD complex are shown.

**Supplementary Figure 2.**
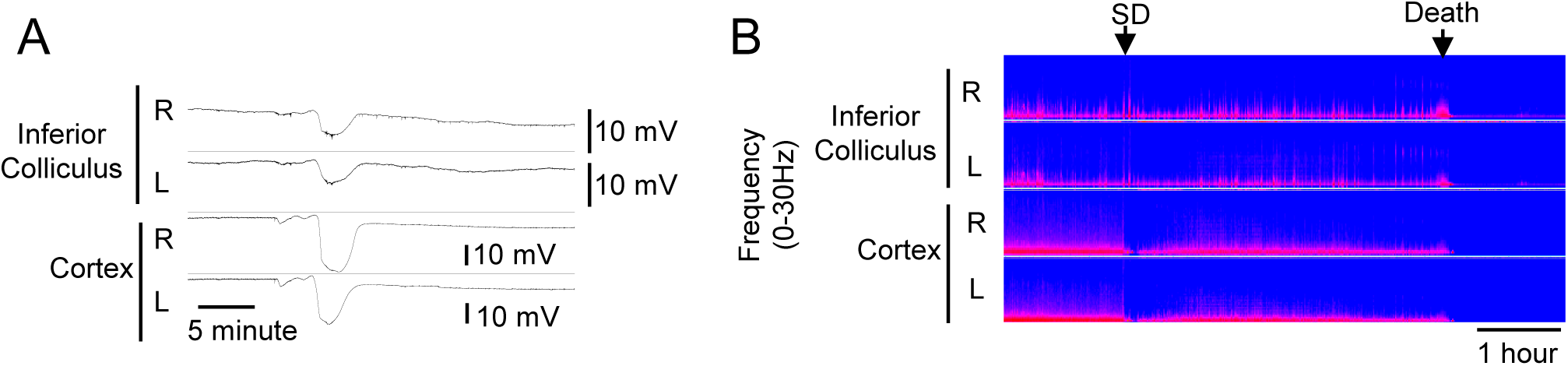
Supplementary data 2. SD in the inferior colliculus in a *Scn8a*^D/+^ mouse. A bilateral SD-like DC potential shift in the inferior colliculus was detected in a *Scn8a*^D/+^ mouse hours before death. **A.** DC traces of SD in right (R) and left (L) inferior and cortical electrodes. **B.** Power spectrum density trace showing the timing of the global depolarization relative to the death.

## Acute slice preparation

Mice were deeply anesthetized with a ketamine/xylazine mixture, cardiac perfused with dissection solution (110 mM NMDG, 26 mM NaHCO_3_, 3 mM KCl, 1.25 mM NaH_2_PO_4_, 6 mM MgSO_4_, 10 mM glucose, 0.4 mM sodium ascorbate, 1 mM thiourea, 0.2 mM CaCl_2_ equilibrated with 95% O_2_/5% CO_2_), and decapitated. The brain was extracted into the ice-cold dissection solution and sectioned (300 µm thick) into either coronal somatosensory slices containing barrel cortex for KCl-SD assays or horizontal cortico-hippocampal slices (anatomical level containing the hippocampal fornix, removing the rostral portion including barrel cortex) for Ca^2+^ imaging. We omitted the barrel cortex from horizontal slices because otherwise our slices could variably contain this SD-susceptible structure, which could result in variable Ca^2+^ responses, even when slices were prepared from the same brain. Once cut, slices recovered for 5 minutes in the dissection solution at 35°C and were maintained in ACSF (126 mM NaCl, 26 mM NaHCO_3_, 3 mM KCl, 2 mM CaCl_2_, 1.25 mM NaH_2_PO_4_, 1 mM MgSO_4_, 0.4 mM sodium ascorbate, 10 mM glucose, equilibrated with 95% O_2_/ 5% CO_2_) at room temperature (22-24°C).

